# Modern human alleles differentially regulate gene expression across brain regions: implications for brain evolution

**DOI:** 10.1101/771816

**Authors:** Alejandro Andirkó, Cedric Boeckx

## Abstract

The availability of high-coverage genomes of our extinct relatives, the Neanderthals and Denisovans, and the emergence of large, tissue-specific databases of modern human genetic variation, offer the possibility of probing the evolutionary trajectory of heterogenous structures of great interest, such as the brain. Using the GTEx cis-eQTL dataset and an extended catalog of *Homo sapiens*-specific alleles relative to Neanderthals and Denisovans, we generated a dataset of nearly fixed, *Homo sapiens*-derived alleles that affect the regulation of gene expression across 15 brain (and brain related) structures. The list of variants obtained reveals enrichments in regions of the modern human genome showing putative signals of positive selection relative to archaic humans, and bring out the highly derived status of the cerebellum. Additionally, we complement previous literature on the expression effects of ancestral alleles in the *Homo sapiens* brain by pointing at a downregulation bias caused by linkage disequilibrium.

## 1 Introduction

State-of-the-art geometric morphometric analyses of endocasts have revealed significant differences between Neanderthal and *Homo sapiens* neurocrania, and have led to the conclusion that specific brain regions, particularly the cerebellum, the parietal and temporal lobes have expanded in the modern lineage as a result of differential growth of neural tissue, with potential consequences for the evolution of modern human cognition [1, 2, 3, 4, 5]. Other differences, affecting subcortical regions that do not leave a direct impact on skulls, are harder to detect, but may also exist [6].

Probing the nature of brain tissue differences is challenging, but the availability of several high-quality archaic genomes [7, 8, 9] has opened numerous research avenues and opportunities to studying the evolution of the *Homo sapiens* brain with unprecedented precision. Early efforts tried to determine the molecular basis of archaic-modern differences based on the few missense mutations that are *Homo sapiens*-specific [10]. But evidence is rapidly emerging in favor of an important evolutionary role of regulatory variants, as originally proposed more than four decades ago [11]. For instance, selective sweep scans to detect areas of the genome under putative positive selection after the split with the Neanderthal lineage show that regulatory variants played a prominent role in ancient selection events [12]. Likewise, changes in potential regulatory elements have been singled out in attempts to identify the factors that gave the modern human face its shape [13, 14]. Other approaches that exploit data from large biobanks have also stressed differences in gene regulatory architecture between modern humans and archaic hominins [15].

Researchers have also explored the idea of connecting genetic variation in modern human genomes, genetic expression analysis and brain evolution. A major study [16] explored the effects of Neanderthal and Denisovan introgressed variants in 44 tissues in modern humans. The authors found downregulation by introgressed alleles in the brain, particularly in the cerebellum and the striatum. In a similar vein, another study [6] examined the effects of archaic introgression on brain and skull shape variability to determine which variants are associ ated with the globularized brain and skull that is characteristic of anatomically modern humans. Here too, the variants with the most salient effects were those found to affect the structure of the cerebellum and the striatum.

We show that derived alleles and genetic regulation data can be used as a complementary source of information about the evolution of the brain. To this end, we took advantage of the data generated in a recent systematic catalog of human genetic variation [17]. This dataset provides an exhaustive collection of derived, *Homo sapiens*-specific alleles found in the present human genetic pool. We chose variants found at a high frequency cutoff (≥90%), and probed the effect of the modern alleles on gene expression compared to ancestral alleles found at low frequencies in modern human genomes.

To determine the predicted effect on gene expression of these derived, modern human-specific alleles, we took advantage of the GTEX database (version 8). By offering information about Expression Quantitative Trait Loci (cis-eQTL) across tissues, the GTEx database forces us to think beyond variants that affect the structure and function of proteins and consider those that regulate gene expression. The GTEx data for the following central nervous system tissues (‘regions’): Amygdala, Caudate, Brodmann Area (BA) 9, BA24, Cerebellum, Cerebellar Hemisphere, Cortex, Hippocampus, Hypothalamus, Nucleus Accumbens, Pituitary, Putamen, Spinal Cord, and Substantia Nigra. Of these samples, Cerebellar Hemisphere and Cerebellum, as well as Cortex and BA9, are to be treated as duplicates [18]. Though not a brain tissue *per se*, the Adrenal Gland was included in our study because of its role in the Hypothalamic-pituitary-adrenal (HPA) axis, an important regulator of the neuroendocrine system that affects behavior.

We wish to stress that our focus on brain (and brain-related) structures in no way is intended as a claim that the brain is the most derived structure in *Homo sapiens* relative to extinct human species. While other tissues (such as the bone structure of the face [19]) undoubtedly display derived characteristics, we have concentrated on the brain in this study because our primary interest lies in cognition and behavior, which is most directly affected by brain-related changes.

Our study contributes to the emerging literature on the evolution of the *Homo sapiens* brain, and highlights novel regulatory changes that deserve further exploration. We show that regions under putative positive selection are enriched in derived, high-frequency (HF) eQTL, reinforcing the important role genetic regulation in human evolution highlighted by previous studies [14, 15, 20]. Our data also complements previous work [16], but differs in an important way: while McCoy and colleagues found a significant downregulation of archaic human alleles in the brain, we only find this effect when not controlling for linkage disequilibrium. Finally, we provide evidence that eQTL affect genetic expression in the cerebellum more than expected by chance, after accounting for effects such as tissue sample size. Additionally, genes affected by eQTL exclusively in the cerebellum are enriched in microtubule-related terms in a GO analysis, suggesting an effect of derived eQTL on cerebellar morphology and development.

## 2 Results

We extracted variation data from [17], a dataset that determines *Homo sapiens* allele specificity using three high-coverage archaic human genomes available at the moment (the Altai and Vindija Neanderthals [7, 8], and a Denisovan individual [9]). The original study [17] introduced an allele frequency cutoff of ≥90% to generate their High-Frequency data subset. We adopted the same filter here, but, departing from the original article data, we decided to restrict our attention to those derived alleles found at ≥90% not only globally, but in each of the major human populations (see Methods section).

The variation data was crossed with the list of variants obtained with the GTEX significant cis-eQTL variants dataset to determine if the selected variants affect gene expression, focusing on 15 central nervous system-related tissues. The GTEx data consist of statistically significant allele effects on gene expression dosage in single tissues, obtained from brain samples of adult individuals aged 20 to 60 [18].

The resulting dataset is composed of *Homo sapiens* derived alleles at high frequency that have a statistically significant effect (at a FDR threshold of 0.05, as defined by the GTEX consortium [21]) on gene expression in any of the selected adult human tissues. In quantitative terms, this amounts to 8,271 statistically significant SNPs associated with the regulation of a total of 896 eGenes (i.e., genes affected by cis-regulation). When controlling for total eQTL variance between brain regions a Chi-square test reveals that the proportion of derived, HF eQTL across tissues is significantly different compared to the rest of non-derived, non-high-frequency eQTL (*p* < 2.2*e* − 16). A post-hoc residual analysis indicates that regions such as the pituitary and the cerebellum are among the major contributors to reject the null hypothesis that the distribution is similar between both groups (*p* < 0.05).

Intronic variants constitute the most abundant category among the derived HF eQTL dataset, but the distribution of categories likely reflects the most common genetic functions near transcription start sites. We controlled for this effect by testing if the functional categories of derived eQTL at high frequency are significantly different from the categories of the rest of GTEx eQTL variants in brain tissues, and found this to be the case (Chi-square test, *p* < 2.2*e* − 16). NMD transcript, non coding transcript, and 5’-UTR variants are the categories driving significance (*p* =< 2.2*e −* 16 for the three sets, residual analysis).

### 2.1 Clumping

To account for linkage disequilibrium and ensure statistical independence, variant clumping was applied through the eQTL mapping p-value at a *r*2 = 0.1. After clumping, the dataset was reduced to 1,270 alleles across tissues, out of which 211 are region-specific (Figure 1B). Because eQTL discovery is highly dependent on the number of tissue samples [21], tissues with more samples tend to yield a higher number of significant variants, regardless of tissue specificity (Figure 1C), as shown by a Spearman correlation test (*p* = 0.0017; *r* = 0.74, controlled for linkage disequilibrium). However, a polynomial regression line fit (blue line in Figure 1C) shows that the cerebellum, adrenal gland and BA9 fall outside the regression’s standard error confidence intervals (in gray in Figure 1C).

**Figure 1:**
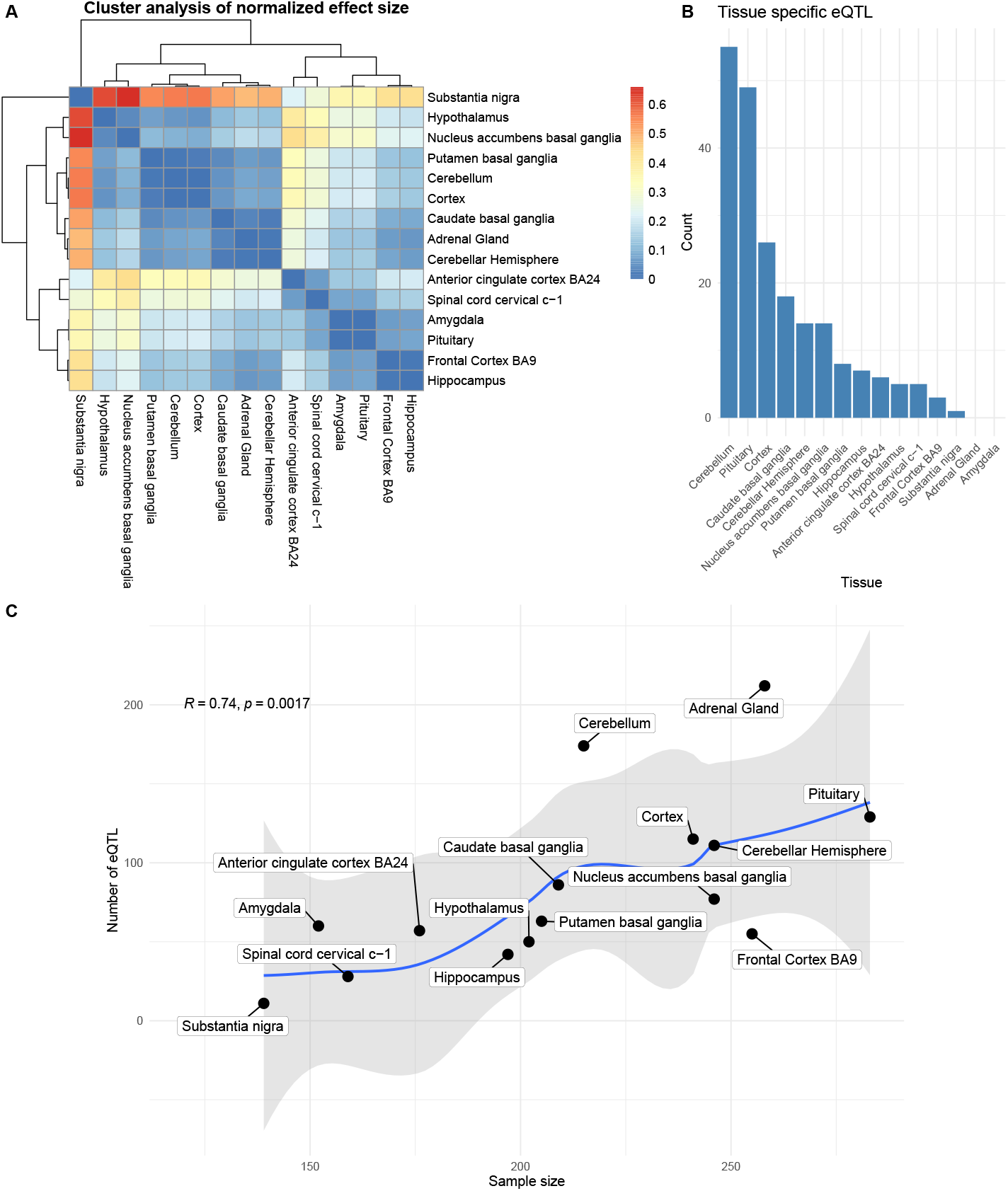
A: Hierarchical clustering analysis of eQTL normal effect size, not controlled for linkage disequilibrium (LD). Color denotes hierarchical distance. B: Number of tissue-specific eQTL after clumping. Adrenal gland and Amygdala do not contain tissue-specific eQTL in our dataset. C: Brain region sample size and eQTL count correlate in our dataset. The blue line marks a polynomial regression line fit, with regression’s standard error confidence intervals (95%) in gray.

We sought to understand if the brain regions just highlighted still stand out considering that most eQTL are shared among regions. The distribution of clumped region-specific variants (Figure 1B) does not correlate with GTEx RNAseq sample size (*p* = 0.9495, Pearson correlation test). The lack of correlation of the region-specific variants with RNAseq sample size might be explained by known effects of genetic regulation disparity between brain regions, such as the distinctive profile of cerebellar eQTL [22, 23].

Additionally, we designed a random sampling testing approach (n=100) to see if any particular region tends to draw more clumped unique eQTL regardless of total eQTL values. The test reveals no significant difference in proportions (*p* = 0.3647, Chi-square independence test). The fact that the adrenal gland and the amygdala have no unique clumped variants might be driving this result.

### 2.2 Directionality of regulation

A previous study [16] had suggested that Neanderthal alleles present in the the modern human genetic pool downregulate gene expression in brain tissue. There is no significant deviance from the expected 50% proportion between down and upregulating variants (*p* = 0.3656, Chi-square test) in our derived HF eQTL dataset (Figure 2B). A significant deviance from the expected 50% proportion (*p* < 2.2*e* − 16, Chi-square test) only obtains when linkage disequilibrium was not controlled for (Figure 2A). A hierarchical cluster analysis of the distance of normalized effect size between regions in non-clumped eQTL shows how the substantia nigra is particularly affected by the downregulating direction skewness effect (Figure 1A).

**Figure 2:**
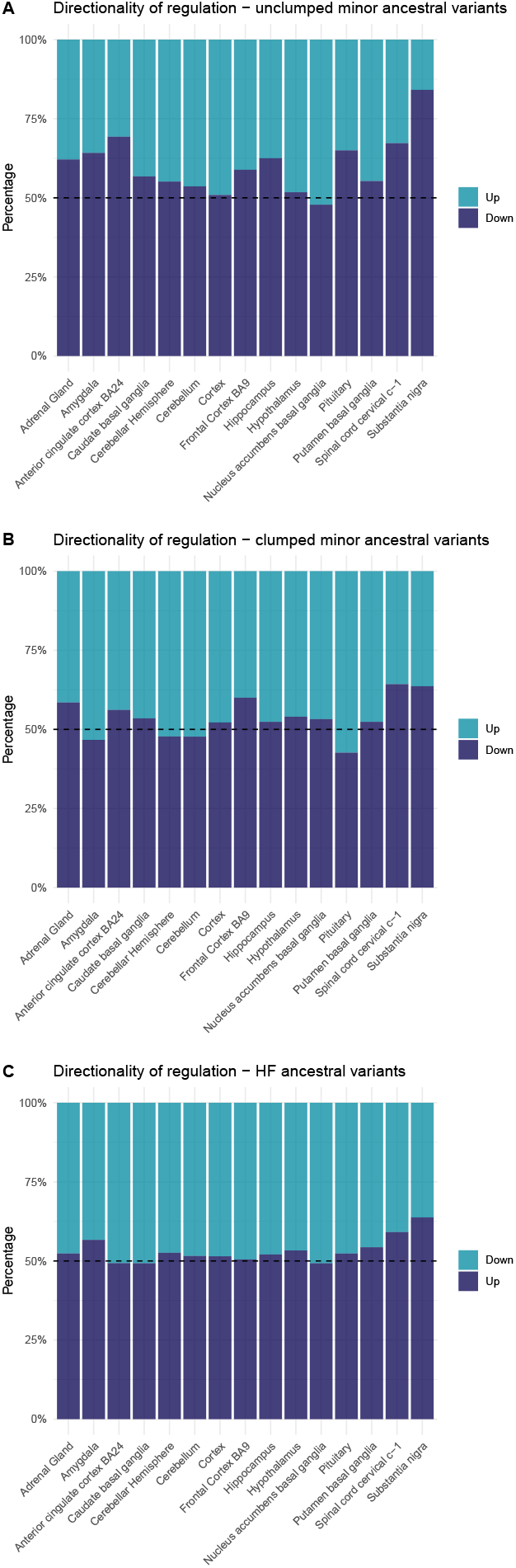
Distribution of up and down-regulating ancestral variants across different subsets of the data, in all eGenes. We include here data before (A) and after (B) controlling for linkage disequilibrium in minor alleles (≥10% frequency). A control using major ancestral alleles (at ≥90% frequency) is included (C).

The same deviation from the expected 50% up and down-regulation proportion was present in major ancestral alleles at a 90% frequency threshold (*p* =< 2.2*e* − 16, Chi-square test, Figure 2C), discarding the possibility that the asymmetry is due to allele frequency cutoffs. However, post-hoc residual analysis shows that downregulating eQTL skewness affects different tissues in the major and minor ancestral eQTL sets. Our analysis suggests that the asymmetric directionality of eQTL regulation is not particular of a given tissue nor is accounted for by frequency. Rather, it appears to be an artifact of failure to take linkage disequilibrium into account.

### 2.3 Regions of evolutionary significance

To determine further the evolutionary significance of any of the variants in our data, we ran two randomization and permutation tests (*N* = 1, 000) to test whether the derived HF eQTL fell within regions under putative positive selection relative to archaic humans as identified in two selective sweep studies ([12, 24]).

We found a significant (*p* = 0.001, observed = 525 overlapping regions, expected = 53) overlap between eQTL and regions of positive selection as defined by [12], as well as in an earlier independent study [24] (*p* < 0.02, observed = 673, expected = 177, Figure 3A and 3B). A Wilcoxon signed-rank test shows that the number of eQTL found in positive selection regions (visualized per region in Figure 3C) is significantly different between studies (*p* = 6.104*e* − 05, after controlling for length differences in the windows detected by each study). A Dunn test (after Bonferroni group correction) failed to find a significant difference between the count of alleles per region in each selective sweep, despite the apparent concordance of the studies in cerebellum (Figure 3C).

**Figure 3:**
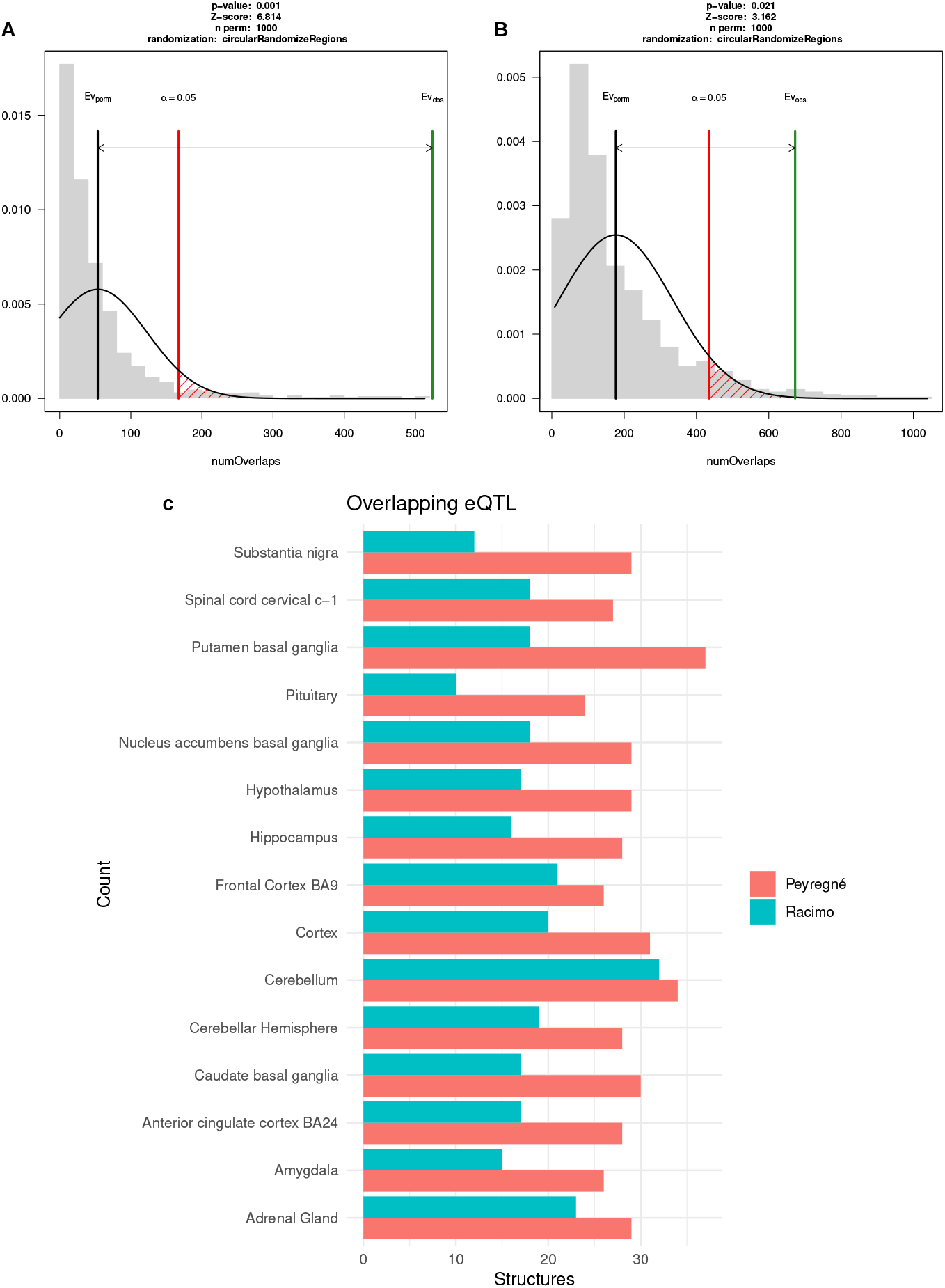
Derived, HF eQTL are present more than expected by chance in selective sweeps from [12] (A) and [24] (B). C shows the count of eQTL overlapping with regions under putative positive selection per region.

Additionally, we tested whether any of the eQTL fell within deserts of introgression, i.e., genetic windows of at least 10 Mb in the *Homo sapiens* genetic pool that have resisted genetic flow from Neanderthals and Denisovans ([25]). While some eQTL do fall within these regions, a permutation test showed that deserts of introgression are not significantly enriched for such variants (*p* > 0.18). We also explored whether derived eQTL overlapped with any known human miRNA or miRNA seeds (as defined in [26]), but found no overlap with our data.

Finally, we tested whether any of the brain-related eQTL were found in genomic locations with a high score according to Pybus et al.’s ([27]) implementation of Fay and Wu’s H test of positive selection [28]. Fay and Wu’s H test doesn’t require ancestral sequences to detect selective sweeps, circumventing the low number of archaic genomes at our disposal. However, derived HF eQTL don’t lie withing regions given a high score by Fay and Wu’s H test in CEU, CHB or YRI populations (at a FDR threshold of 0.01).

### 2.4 Enrichment analysis

A GO enrichment analysis for the clumped variants in the 15 regions we focused on here revealed an over-representation of the categories ‘cytoplasm’, ‘catalytic activity’, and ‘ion binding’ (*p* < 0.05).

Given the importance of the cerebellum in previous studies of great relevance here [1, 6, 16], and the overrepresentation of that region in our eQTL dataset, we ran a GO analysis for cerebellum-affecting eQTL. We found that these derived HF eQTL lie on genes involved in microtubule-related functions (GO categories ‘microtubule binding’ and ‘microtubule binding’). Microtubules play an important role in cerebellar neuronal migration and in maintaining morphological stability through development [29].

In an attempt to link the derived eQTL to phenotypical effects as detected by GWAS, a phenome-wide association (PheWAS) query [30] was run on the clumped dataset of variants. Several variants are top hits in GWAS related to the immune system and other traits of interest, such as bone mineral density, brain volume in various regions and lymphocyte cell count. However, a downstream colocalization analysis did not find significant results in any of the selected GWAS (including for traits previously claimed to be derived in human evolution studies, such as cerebellar volume or Alzheimer’s disease [6, 31]).

## 3 Discussion

In this study we sought to shed light on the contribution of modern-human-specific alleles found at high frequency in differential gene regulation across brain regions. In so doing we hoped to complement previous work that focused on the effects of introgressed variants [6, 16], as well as provide an alternative approach to studying fixed variants exclusively [10]. We have shown that the cerebellum accumulates more derived HF eQTL than expected by chance, supporting previous claims about the derived nature of the cerebellum in the context of modern human evolution [3, 6, 13].

We did not find a significant skewness towards downregulation in derived eQTL, regardless of frequency. This effect was previously detected as a characteristic of Neanderthal alleles introgressed in the modern human genetic pool [16]. The derived eQTL did show directional regulatory asymmetry but only when linkage disequilibrium was not controlled for. Additional testing indicates that the effect is not introduced by the high frequency cutoff imposed to the data, nor introduced by the bias of a particular region in either HF or non-HF alleles.

We also found that regions of putative positive selection exhibit an enrichment for derived eQTL. The authors of [12] introduced fixed alleles as an additional source for their selective sweeps, accounting for a mean difference of 3% minor allele frequency in our eQTL dataset compared to the putative positive selection alleles (as per frequency values reported in [12]’s supplementary file 3). This difference in minor allele frequency might affect the number of detected eQTL in regions of positive selection, as the detection power of eQTL negatively correlates with MAF [18], making near-fixation alleles harder to map as eQTL. We suggest that derived HF eQTL might affect the modern human genetic regulation landscape by either being drivers of positive selection or being in linkage disequilibrium with causal positively selected variants. This is in agreement with the authors of one of the selective sweep studies, who found that regions under putative positive selection are enriched in regulatory variants [12].

Some of the genes associated with signals of positive selection and affected by differential gene expression have already been linked to clinical phenotypes or brain development. For example, *NRG4* is involved in dendritic development [32], RAB7A has been found to be related to tau secretion, a marker of Alzheimer’s disease [33], a disease hypothesized to be human-specific [31], and GABPB2 has been associated with schizophrenia [34]. We highlight as well a derived eQTL in the BAZ1B gene that lies in one of the regions under putative positive selection. This variant affects the expression of two genes in cerebellar tissue that, like *BAZ1B* itself, are part of the Williams-Beuren Syndrome Critical Region (*MLXIPL* and *NSUN5P2*). BAZ1B is known to be related to craniofacial development in human evolution [13].

All in all, our work reinforces the potential of using human variation databases as a valuable point of entry to connect genotype and phenotype in brain evolution studies, and corroborates claims about the importance of genetic regulation in human brain evolution.

## 4 Methods

We accessed the *Homo sapiens* variant annotation data from [17]. The original complete dataset is publicly available at https://doi.org/10.6084/m9.figshare.8184038. This dataset includes archaic-specific variants and all loci showing variation within modern populations, using the 1000 genomes project and ExAc data to derive frequencies and the human genome version *hg19* as reference. As described in the original article, the authors also applied quality filters in the archaic genomes (sites with less 5-fold coverage and more than 105-fold coverage for the Altai individual, or 75-fold coverage for the rest of archaic individuals were filtered out). In ambiguous cases, variant ancestrality was determined using multiple genome aligments [35] and the macaque reference sequence (*rheMac3*) [36].

For replication purposes, we wrote a script that reproduces the 90% frequency cutoff point used in the original study. We filtered the variants according to the guidelines in [17] such that: 1) all variants show 90% allele frequency, 2) the major allele present in *Homo sapiens* is derived (ancestrality is either determined by the criteria in [35] or by the macaque reference allele), whereas either archaic reliable genotypes have the ancestral allele, or the Denisovan carries the ancestral allele and one of the Neanderthals the derived allele (accounting for gene flow from *Homo sapiens* to Neanderthal).

Additionally, the original study we relied on [17] applies the 90% frequency cutoff point in a global manner: it requires that the global frequency of an allele be more than or equal to 90%, allowing for specific populations to display lower frequencies. Using the metapopulation frequency information provided in the original study, we applied a more stringent filter and removed any alleles that where below 90% in any of the five major metapopulations included (African, American, East Asian, European, South Asian). We then harmonized and mapped the high-frequency variants to the data provided by the GTEx database [21]. In order to do so we pruned out the alleles that did not have an assigned rsIDs.

Post-mostem mRNA degradation affects the number of discovered eQTL in other tissues. However, we did not control for post-mortem RNA degradation, since the Central Nervous System has been shown to be relatively resistant to this effect. [37]. However, re-sampled tissues (here labeled ‘cerebellar hemisphere’ and ‘Cortex’ following the original GTEx Consortium denominations) do show differences compared to their original samples (‘cerebellum’ and ‘BA 9’). We acknowledge that the resulting data are limited by inherent problems of the GTEx database, such the use of the same individuals for different brain tissue samples, the reduced discovery power of rare variants [18] or artifacts introduced during RNAseq analysis.

Clumping of the variants to control for Linkage Disequilibrium was done with Plink (version 1.9) through the *ieugwasr* R package [30], requiring a linkage disequilibrium score of 0.90 (i.e., co-inheritance in 90% of cases) for an SNP to be clumped. The nominal p-value of eQTL mapping was used as the criterion to define a top variant; i.e., haplotypes were clumped around the most robust eQTL candidate variant. Linkage disequilibrium values are extracted from the 1000 Genomes project ftp server (ftp://ftp.1000genomes.ebi.ac.uk/vol1/ftp/release/20130502/) by the *ieugwasr* R package.

Distance values for tissue hierarchical clustering were calculated by using the mean values of the normalized effect size of derived HF eQTL.

We performed the permutation test (n=1,000) with the R package RegioneR using the unclumped data, as variants might clump around an eQTL falling outside windows of putative positive selection, underepresenting the number of data points inside such genomic areas and reducing statistical power.

We performed the Gene Ontology analysis with the *gprofiler* R package [39], using as background the whole genome, at a *p* = 0.05 significance threshold. We performed the Phenome-wise association scan (PheWAS) (at a *p* = 0.0001 threshold) and colocalization analysis (at a *p* = 5*e* − 04 threshold for top hit identification) through the *ieugwasr* [30], *MRinstruments* and *gwasglue* packages. The selected GWAS for colocalization can be consulted in the relevant section of the article’s code.

Figures were created with the ggplot2 R package [40] and RegioneR [38]. All statistical tests were controlled for power (≥ 0.8). The complete code to reproduce the data processing, plot generation and analysis can be found in https://github.com/AGMAndirko/GTEX-code. The miRNA data was extracted from the Supplementary Tables S6 and S7 of [26]. The human selective sweep data was extracted Supplementary Table S5 from [24], and Supplementary Table S2 from [12]. For the deserts of introgression data we extracted the information from [25].

## Acknowledgments

The Genotype-Tissue Expression (GTEx) Project was supported by the Common Fund of the Office of the Director of the National Institutes of Health, and by NCI, NHGRI, NHLBI, NIDA, NIMH, and NINDS. The data used for the analyses described in this manuscript were obtained from the GTEx Portal on 05/15/19.

## Author Contributions

Conceptualization: CB & AA; Data Curation: AA; Formal Analysis: AA; Funding Acquisition: CB; Investigation: CB & AA; Methodology: CB & AA; Software: AA; Supervision: CB; Visualization: CB & AA; Writing — Original Draft Preparation: CB & AA; Writing — Review & Editing: CB & AA.

